# Vascular risk factors for male and female urgency urinary incontinence at age 68 from a British birth cohort study

**DOI:** 10.1101/246462

**Authors:** Alex Tsui, Diana Kuh, Linda Cardozo, Daniel Davis

**Author notes:** Corresponding author: Alex Tsui, MRC Unit for Lifelong Health and Ageing at UCL, 33 Bedford Place, London, UK, WC1B 5JU.

## Abstract

**Objective:** To investigate the prevalence of UUI at age 68 and the contribution of vascular risk factors to male and female UUI pathogenesis in addition to the associations with raised BMI

**Subjects and methods:** 1762 participants were from the MRC National Survey for Health and Development (NSHD) birth cohort, who answered the International Consultation on Incontinence Questionnaire short form (ICIQ-SF) at age 68. Logistic regression was used to estimate associations between UUI and earlier life vascular risk factors including: lipid status, diabetes, hypertension, body mass index (BMI), previous stroke or transient ischaemic attack (TIA) diagnosis; adjusting for smoking status, physical activity, co-presentation of SUI symptoms, educational attainment and in women only, type of menopause, age at period cessation and use of hormone replacement therapy.

**Results:** UUI was reported by 12% of men and 19% of women at 68. Female sex, previous stroke or TIA diagnosis, increased BMI and hypertension (in men only) at age 60-64 were independent risk factors for UUI. Female sex, increased BMI and a previous diagnosis of stroke/ TIA increased the relative risk of more severe UUI symptoms. Type and timing of menopause and HRT use did not alter the estimated associations between UUI and vascular risk factors in women.

**Conclusion:** Multifactorial mechanisms lead to UUI and vascular risk factors may contribute to pathogenesis of bladder overactivity in addition to higher BMI. Severe UUI appears to be a distinct presentation with more specific contributory mechanisms than milder UUI.

## Introduction

Urgency urinary incontinence (UUI) is involuntary loss of urine associated with urgency (a sudden compelling desire to pass urine which is difficult to defer) (1), commonly presenting with urinary frequency and nocturia as part of the overactive bladder (OAB) syndrome. UUI is common: prevalence of 30% has previously been reported for those over 65 (2, 3). The actual prevalence of UUI is likely to be even higher, as many older people may be too embarrassed to report symptoms (4, 5). Greater frequency or severity of UUI symptoms are associated with worse quality of life (2, 6–8) and increased risk of depression (9), negatively impacting social interactions, relationships and self-esteem (10). The cost of OAB syndrome in the UK alone is currently estimated at £800 million per year, with a further 22% increased anticipated by 2020 as a result of medications, increased need for social care and higher risk of admission to a nursing home (11).

The multifactorial causes of UUI are reflected in the wide range of risk factors identified in cross-sectional and longitudinal studies. These include increasing age (12), diagnosis of depression (13), alcohol intake (14) and limitations in physical capability (15). In addition, increased BMI has been consistently associated with all types of urinary incontinence including UUI in a number of cohorts (16–18). It is understood that central adiposity increases intra-abdominal and bladder pressure in SUI and to a lesser extent UUI (3). However, increased BMI is also commonly associated with vascular risk factors, such as insulin resistance and glucose intolerance, dyslipidaemia and hypertension, as part of a metabolic syndrome (19). As such, vascular mechanisms may contribute to UUI pathophysiology at the peripheral and central nervous systems, as well as directly on the detrusor muscle and pelvic soft tissue.

Although UUI impacts a significant proportion of both sexes in the population, risk factors for UUI have mostly been explored in cross-sectional samples of women (17, 18, 20–22). In this study, we investigated vascular risk factors assessed at age 60-64 and UUI subsequently ascertained in men and women at age 68 within a birth cohort, taking account of a number of clinical and socio-behavioural factors. We addressed four main questions:

1. What is the prevalence of male and female urinary incontinence subtypes at age 68?
2. Do vascular risk factors contribute to male and female UUI in addition to the effects of raised BMI?
3. Is there evidence for a central neurological contribution to UUI, through the additional contribution of a stroke/TIA diagnosis over and above vascular risk factors in general?
4. Are the same vascular risk factors observed in association with severe and mild UUI?

## Methods

The MRC National Survey for Health and Development (NSHD) is the oldest British birth cohort study, following a sample of 5362 male and female participants, born in one week in March 1946. During the 24^th^ data collection in 2014-5, 2942 participants in the target sample living in England, Scotland and Wales were contacted and 2453 (83.4%) returned a postal questionnaire.(23) The target sample did not include participants no longer in the study (n=2420): 957 (17.8% of the original sample) had already died, 620 (11.6%) had previously withdrawn from the study, 448 (8.3%) had emigrated and had no contact with the study, and 395 (7.4%) had been untraceable for more than five years. The 2294 participants (93.5% of 2453 who had returned a postal questionnaire), whose symptoms reported on the postal questionnaire corresponded to a recognised incontinence syndrome were used as the maximum sample for analysis. Of these, 1753 (76.4%) men and women had full covariate data (detailed below), and comprised the complete sample (Figure 1).

### Urinary Leakage

Questions on urinary leakage were based on the International Consultation on Incontinence Questionnaire short form (ICIQ-SF) (24): *“1. How often do you leak urine? 2. How much urine do you usually leak? 3. How much does leaking urine interfere with your everyday life, on a scale of 0 to 10?”*

Urinary symptoms were categorised into those with (i) UUI only; (ii) stress urinary incontinence (SUI) only; (iii) mixed incontinence (MUI). Participants with UUI were defined as those who reported urine leakage “before you can get to the toilet” or “when you are asleep”. Participants with SUI were defined by urine leakage “when you cough or sneeze” or “when you are physically active or exercising”. Mixed incontinence was defined as responses with combination of SUI and UUI. A severity score was calculated from the ICIQ-SF using the sum of question 1, question 2 score multiplied by 2 and question 3, as recommended (24). A severity score was not calculated for 14 participants who did not respond to all three components. Incontinence severity was defined as: no incontinence (score of 0), mild incontinence (3–5) and more severe incontinence (>=6).

### Vascular exposures

At age 60-64, information was collected by a research nurse at a home or clinic visit (25) on vascular risk factors typically recognised as a component of the metabolic syndrome (19): lipid status, diabetes, hypertension, body mass index (BMI) and waist circumference. Hypertension was defined as a doctor diagnosis of hypertension, regular prescription of an anti-hypertensive, or systolic blood pressure of greater than 160 mmHg or diastolic blood pressure greater than 100 mmHg (taken from two readings) (26). Participants reported doctor-diagnosed type 1 or 2 diabetes mellitus. During the home or clinic visits, waist circumference, height and weight were measured by standardised protocols, and BMI was calculated (kg/metres^2^). A fasting blood sample was also taken during this home visit. Lipid status was defined according to whether the participant was hypercholesterolaemic (total cholesterol >6mmol/L) and/or if a cholesterol-lowering medication was prescribed. At age 68, participants reported any previous diagnosis of stroke/ transient ischaemic attack (TIA) ascertained by a doctor.

### Other covariates

Other covariates selected included: smoking status (defined as current smoker, ex-smoker, or lifelong non-smoker, validated against reports at earlier ages); co-presentation of SUI symptoms at age 68; physical activity at age 60-64, and educational attainment by age 26. Educational attainment was categorised into: less than ordinary secondary level; ‘O’ levels; advanced secondary level (‘A’ level) and higher. Participants were asked how many times in the last four weeks they had taken part in sports or vigorous activities, categorised as inactive (no episodes), less active (1-4 exercise episodes per month) and more active (five or more exercise episodes per month). In women, we also accounted for type of menopause (natural menopause, bilateral oophorectomy or hysterectomy with at least one conserved ovary), age at period cessation and ever-use of hormone replacement therapy (27).

### Statistical analysis

First, using the maximum sample, the proportion of men and women with UUI at age 68 by each vascular risk factor and covariate was described. Logistic regression was used to assess the strength of the associations for men and women separately, testing for any sex interactions. Second, we repeated the logistic regressions in 1762 men and women with complete covariate data, and made a series of three adjustments: (1) for female sex, SUI, and previous diagnosis of stroke/ TIA; (2) additionally for vascular risk factors; and (3) additionally for all other covariates. Analyses adjusted for BMI per standard deviation rather than waist circumference as the measure of adiposity; using both would have resulted in collinearity. Third, we estimated multinomial logistic regression models to compare risk factor profiles between those with severe and mild UUI symptoms. Fourth, we estimated logistic regression models to investigate the associations of type and timing of menopause and HRT use on UUI in women to test whether any observed associations between UUI and vascular risk factors could be accounted for by menopause variables.

## Results

Of 2294 participants, 825 reported symptoms at age 68 consistent with a recognised urinary incontinence subtype. For men, 15% reported incontinence (UUI 12%, SUI 1.5%, MUI 1%); for women, 54% reported incontinence (UUI 19% SUI 21% MUI 14%), demonstrating a sex difference (p<0.01). In men, 7% with an incontinence subtype had severe symptoms, compared with 18% of women.

Co-presentation of SUI was associated with UUI in both sexes, but was stronger in men than women (men = OR 5.3, 95% confidence intervals (CI) 2.4 to 11.4; women = OR 1.6 (95% CI 1.2 to 2.0)) (Table 1). Those diagnosed with a previous stroke or TIA reported more UUI (26.3% vs 12.5% for men (odds ratio (OR) 2.6 (95% CI 1.4 to 4.8)); 47.5% vs 32.6% for women, (OR 1.9 (95% CI 1.0 to 3.6)). A diagnosis of hypertension was associated with an increased UUI risk in men and women; diabetes was associated with an increased UUI risk in men but not in women (sex interactions with hypertension p=0.49, with diabetes, p=0.03). Raised BMI was associated with increased UUI risk in men and women. No associations between physical activity and UUI were evident.

**Table 1:**
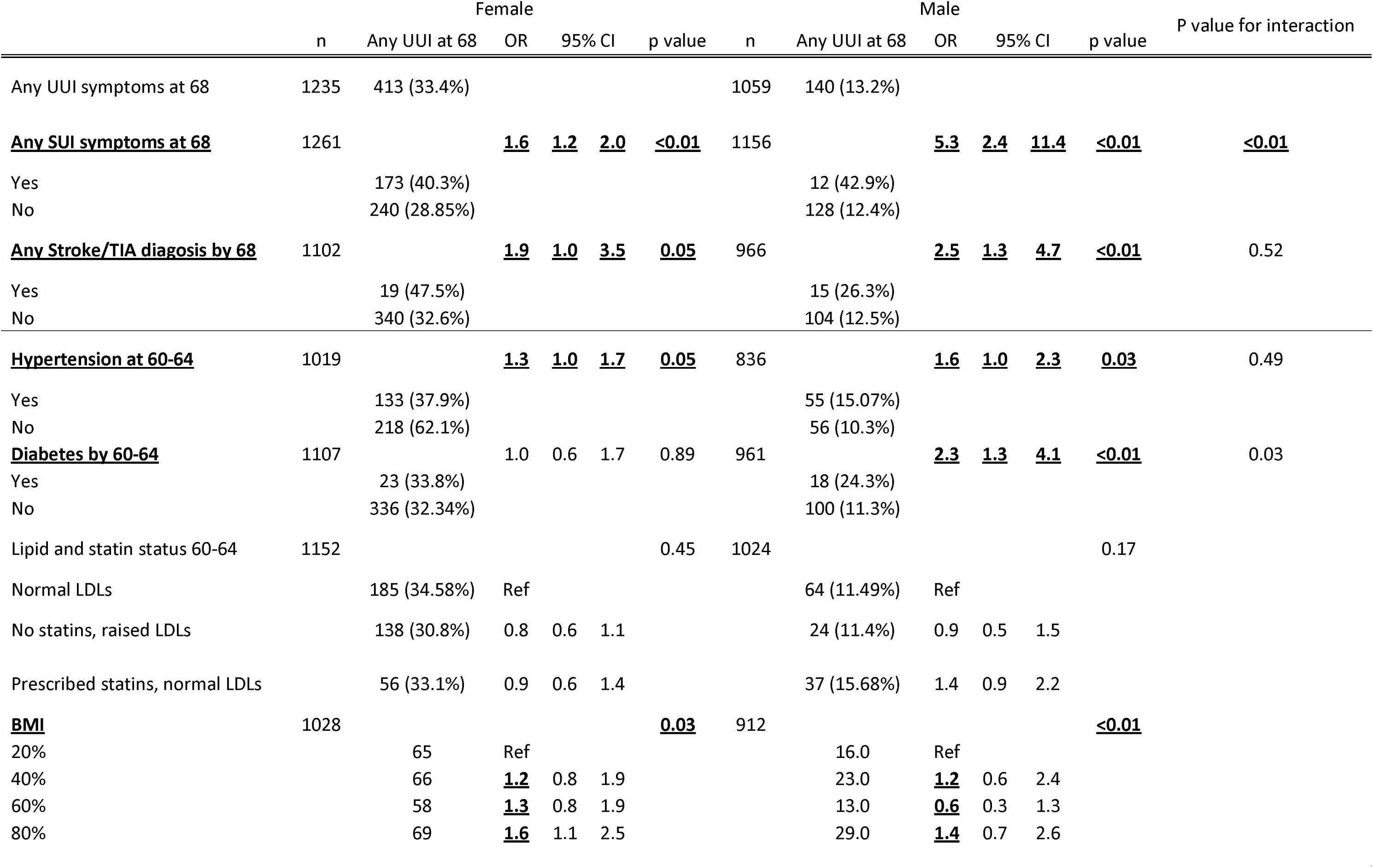
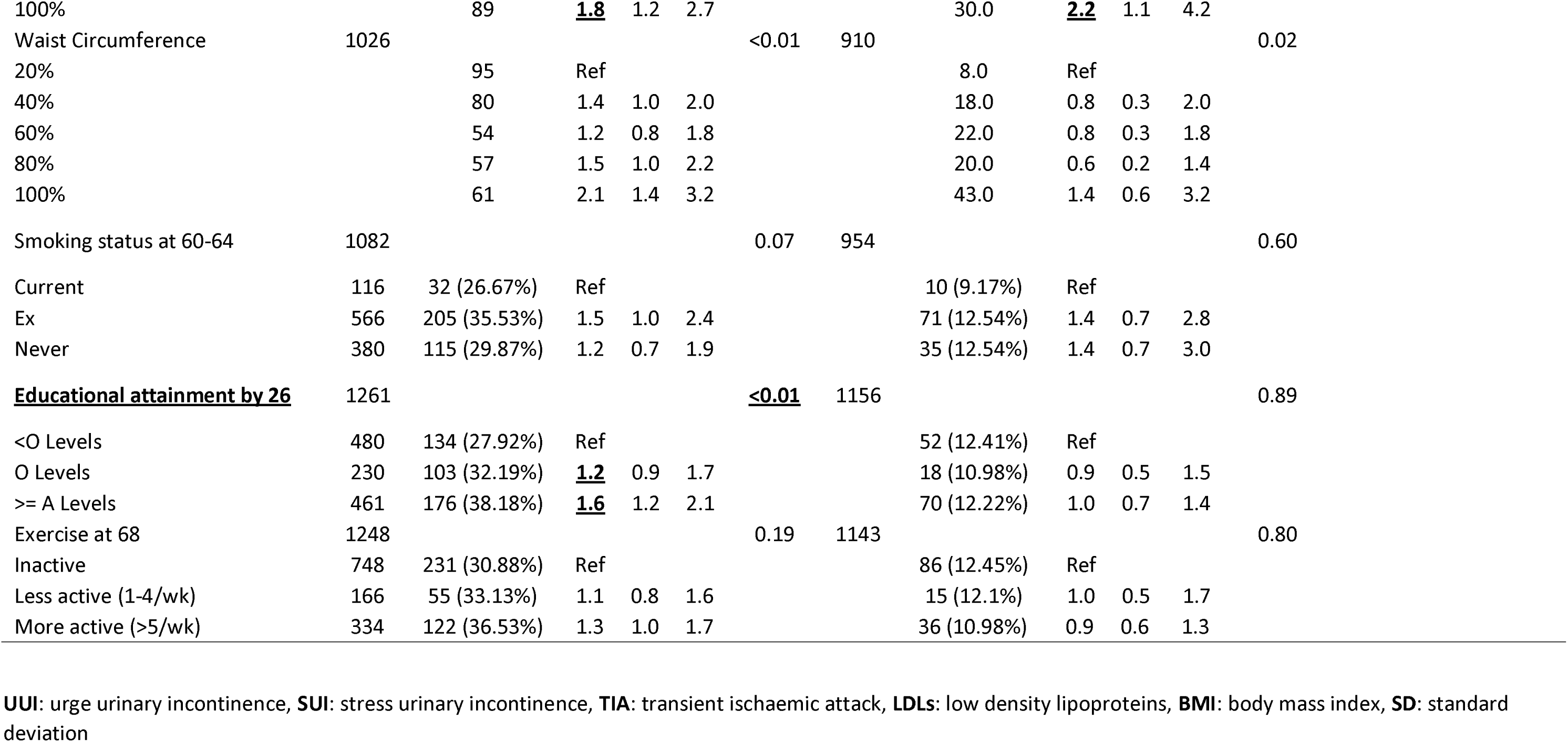
Prevalence of UUI and SUI, and odds ratios for risk factors of UUI in maximum sample of male and female participants in National Survey of Health and Development (NSHD) cohort

Univariate models in those with complete data confirmed that being female, SUI symptoms, a previous stroke or TIA diagnosis, increased BMI, hypertension and diabetes (in men only) continued to be associated with higher odds of UUI (Table 2, model 1). There were no associations between smoking status, educational attainment or physical activity and UUI. After adjusting for sex, SUI and any stroke or TIA diagnosis the odds ratios for the other vascular factors were attenuated but independent associations with BMI and hypertension remained (model 2). In the fully adjusted model, being female (OR 4.12, 95%CI 2.49 to 6.82, p=<0.01), co-presentation of SUI (OR 1.8, 95%CI 1.36 to 2.37, p=<0.01), having had a stroke or TIA diagnosis (OR 1.99, 95%CI 1.14 to 3.49, p<0.01) and increased BMI (OR 1.19 per SD, 95%CI 1.05 to 1.34, p=0.01) were independent risk factors for UUI.

**Table 2:**
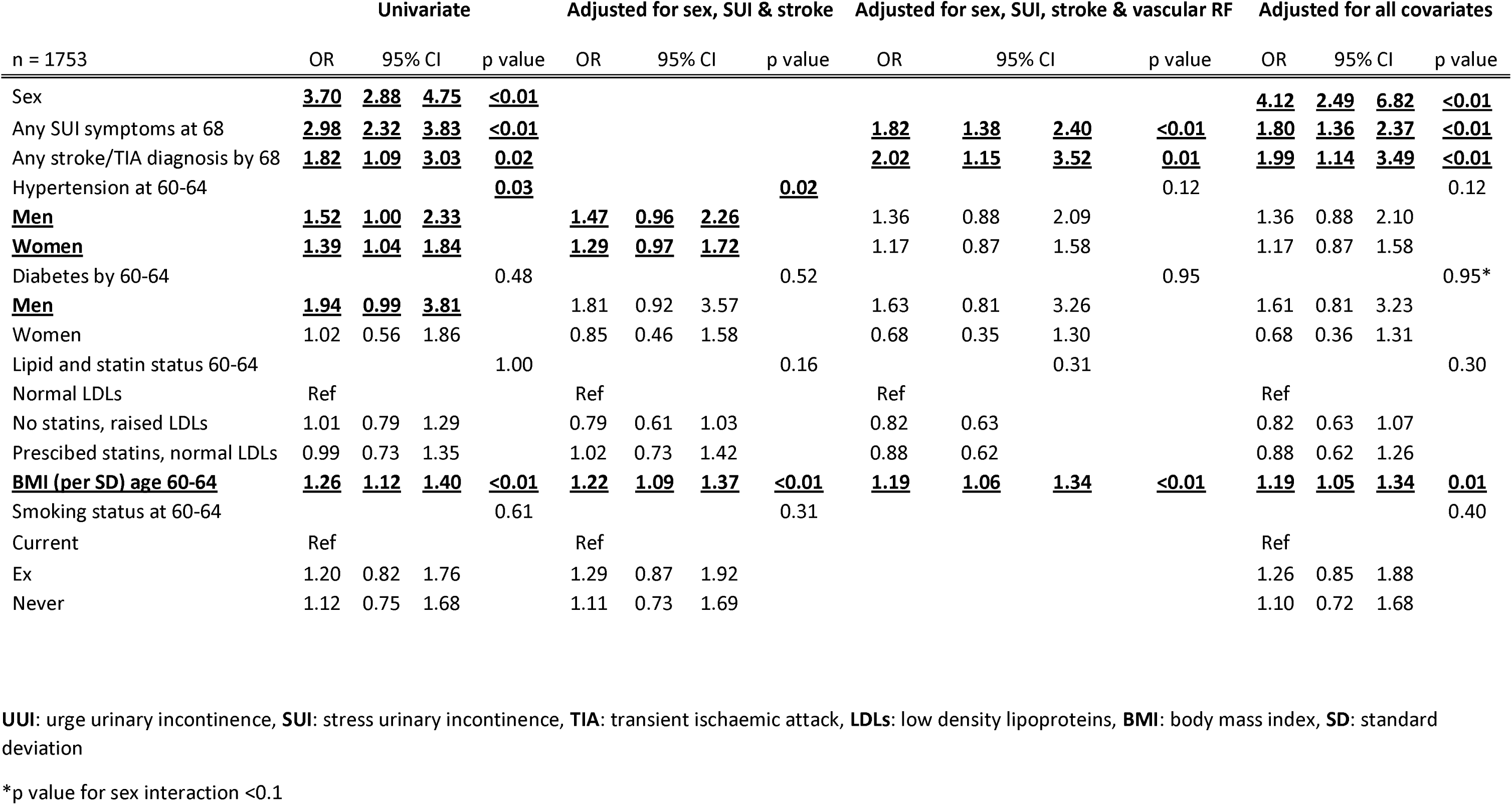
Odds ratios for UUI risk factors in complete sample of 1766 participants from NSHD birth cohort with complete data for vascul ar risk factors

A previous diagnosis of stroke/ TIA increased the relative risk of severe UUI symptoms (RRR 3.65, 95%CI 1.87 to 7.1); no corresponding association was demonstrated with mild UUI. Increased BMI and being female were risk factors for both mild and severe UUI (Table 3). There were no sex interactions in these multivariate models. Type and timing of menopause and HRT use did not alter the estimated associations between UUI and vascular risk factors in women (Table 4).

**Table 3:** Multinomial regression analyses of relative risk ratios in mild and severe UUI in 1761 participants from NSHD with complete da ta for vascular risk factors and incontinence severity scores

| n = 1748 | Adjusted for all covariates |  |  |  |  |  |  |  |
| --- | --- | --- | --- | --- | --- | --- | --- | --- |
|  | Mild UUI |  |  |  | Severe UUI |  |  |  |
|  | RRR | 95% CI |  | p value | RRR | 95% CI |  | p value |
| **<u>Female sex</u>** | **<u>3.51</u>** | **<u>2.58</u>** | **<u>4.77</u>** | **<u><0.01</u>** | **<u>4.96</u>** | **<u>3.25</u>** | **<u>7.58</u>** | **<u><0.01</u>** |
| Any stroke/TIA diagnosis by 68 | 1.08 | 0.49 | 2.40 | 0.84 | **<u>3.65</u>** | **<u>1.87</u>** | **<u>7.10</u>** | **<u><0.01</u>** |
| Hypertension at 60-64 | 1.21 | 0.90 | 1.63 | 0.21 | 1.31 | 0.89 | 1.91 | 0.17 |
| Diabetes by 60-64 | 0.96 | 0.52 | 1.78 | 0.89 | 1.18 | 0.60 | 2.34 | 0.63 |
| Lipid and statin status 60-64 |  |  |  |  |  |  |  |  |
| No statins, normal LDLs | 0.95 | 0.70 | 1.28 | 0.72 |  |  |  |  |
| No statins, raised LDLs | 0.93 | 0.61 | 1.42 | 0.74 | 0.66 | 0.44 | 0.99 | 0.05 |
| Prescribed statins, normal LDLs |  |  |  |  | 0.82 | 0.49 | 1.38 | 0.46 |
| **<u>BMI age 60-64</u>** | **<u>1.13</u>** | **<u>0.98</u>** | **<u>1.30</u>** | **<u>0.09</u>** | **<u>1.32</u>** | **<u>1.12</u>** | **<u>1.57</u>** | **<u><0.01</u>** |
| Smoking status at 60-64 |  |  |  |  |  |  |  |  |
| Current | Ref |  |  |  | Ref |  |  |  |
| Ex | 1.34 | 0.82 | 2.20 | 0.24 | 1.19 | 0.67 | 2.13 | 0.55 |
| Never | 1.37 | 0.82 | 2.29 | 0.23 | 0.71 | 0.37 | 1.35 | 0.30 |

**UUI:** urge urinary incontinence, **SUI:** stress urinary incontinence, **TIA:** transient ischaemic attack, **LDLs:** low density lipoproteins, **BMI:** body mass index, **SD:** standard deviation

**Table 4:** Odds ratios for UUI in female NSHD participants with complete data for women’s health and vascular risk variables

| n=752 | Univariate (complete n) |  |  | Adjusted for all covariates |  |  |
| --- | --- | --- | --- | --- | --- | --- |
|  | OR | 95% CI | p value | OR | 95% CI | p value |
| Type of menopause |  |  |  |  |  |  |
| Natural | Ref |  | 0.29 | Ref |  |  |
| Hysterectomy | 1.25 | 0.83 1.88 |  | 1.65 | 0.94 2.88 | 0.08 |
| Bilateral Oophorectomy | 1.38 | 0.85 2.26 |  | 1.54 | 0.89 2.64 | 0.12 |
| Period cessation age (yrs) |  |  |  |  |  |  |
| Natural | 1.30 | 0.80 2.09 | 0.29 | 1.28 | 0.79 2.08 | 0.32 |
| Hysterectomy | 1.17 | 0.61 2.24 | 0.64 | 1.16 | 0.60 2.25 | 0.67 |
| Bilateral Oophorectomy | 2.09 | 0.81 5.39 | 0.13 | 2.17 | 0.83 5.67 | 0.11 |
| Ever Used HRT |  |  |  |  |  |  |
| Natural | 0.92 | 0.65 1.32 | 0.59 | 0.89 | 0.62 1.28 | 0.53 |
| Hysterectomy | 0.66 | 0.27 1.63 | 0.48 | 0.74 | 0.29 1.88 | 0.53 |
| Bilateral Oophorectomy | 3.66 | 0.41 32.95 | 0.24 | 4.83 | 0.47 49.36 | 0.18 |
| **<u>Any SUI symptoms at 68</u>** | **<u>1.60</u>** | **<u>1.17 2.19</u>** | **<u><0.01</u>** | **<u>1.58</u>** | **<u>1.14 2.17</u>** | **<u>0.01</u>** |
| Any stroke/TIA diagnosis by 68 | 1.79 | 0.78 4.11 | 0.17 | 1.60 | 0.67 3.82 | 0.29 |
| Hypertension at 60-64 | 1.26 | 0.92 1.73 | 0.15 | 1.13 | 0.80 1.59 | 0.48 |
| Diabetes by 60-64 | 1.00 | 0.50 1.98 | 0.99 | 0.76 | 0.35 1.64 | 0.48 |
| Lipid and statin status 60-64 |  |  | 0.37 |  |  | 0.33 |
| Normal LDLs | Ref |  |  | Ref |  |  |
| No statins, raised LDLs | 0.79 | 0.57 1.10 |  | 0.81 | 0.58 1.13 |  |
| Prescribed statins, normal LDLs | 0.86 | 0.54 1.38 |  | 0.74 | 0.44 1.25 |  |
| **<u>BMI age 60-64</u>** | **<u>1.17</u>** | **<u>1.02 1.34</u>** | **<u>0.02</u>** | 1.14 | 0.99 1.33 | 0.07 |

UUI: urge urinary incontinence, SUI: stress urinary incontinence, TIA: transient ischaemic attack, LDLs: low density lipoproteins, BMI: body mass index, SD: standard deviation.

## Discussion

In a large representative British population cohort at age 68, the prevalence of urinary incontinence was 15% in men (where UUI was the most common subtype), and 54% in women (with similar proportions of UUI and SUI subtypes). Female sex, a previous diagnosis of stroke/ TIA, increased BMI and SUI were associated with UUI symptoms. Stronger associations were found between these risk factors and UUI if severe UUI was reported. For those with milder symptoms, the associations were weaker, except for the negative association with educational attainment. In women, no associations were found between UUI symptoms and menopause or HRT use. Taken together, these results suggest that vascular risk factors, in particular hypertension, may contribute towards UUI pathophysiology in addition to previous stroke/TIA, raised BMI, female sex and co-presentation with SUI.

A strength of these analyses was the prospective ascertainment of all variables. We used the ICIQ, a structured, validated scale with a specific component evaluating impact on daily life. A limitation is the lack a definite operationalisation of incontinence subtypes in ICIQ, excluding a small number of participants reporting only atypical symptoms. Also, the severity of incontinence was measured using self-reported symptom impact, without objective urodynamic measures. Third, vascular covariates could not be further characterised by cumulative exposure, such as duration since first diagnosis, introducing a dose-dependent element to the associations. Lastly, the definition of previous TIA/stroke did not differentiate between temporary or permanent neurological deficits, size of lesion or neuroanatomical location of stroke.

While the estimated prevalence of female UUI at 68 was similar to other studies with samples of similar ages (21, 22), UUI prevalence in men was higher than that reported (11.7% for men over 65, 95% CI: 9.27-14.14%) in a recent pooled analysis of men over 65 years (28). The increased prevalence of UUI in both men and women from 53 to 68 is consistent with previous single and multi-centre cross-sectional studies reporting increasing age as a major risk factor for UUI (15, 22, 29). Prevalence of vascular risk factors in NSHD were comparable to published literature: the prevalence of diabetes and hypertension at 60-64 were 6.9% and 38% respectively. In addition, 4.8% of participants had been diagnosed with a stroke or reported TIA symptoms by age 68. These are comparable to published literature: the prevalence estimates of hypertension between 55 and 64 years was 45% according to the Health Survey for England statistics report from 2012 (30); 6.5% for diabetes at ages 55-64 years (31); and 3% for men and 2% for women for prevalence of stroke between 55-64 using the British Heart Foundation stroke statistics 2009 (32). The remaining discrepancies are likely explained by the 100% white Caucasian demographics of the NSHD cohort, while other analyses included participants of Afro-Caribbean and South Asian ethnicities, where prevalence of hypertension and diabetes respective are generally higher.

Our findings suggest severe UUI to be a distinct entity compared to mild UUI at this age, with more specific risk factors. First, a syndrome of mild UUI possibly reflects increased recognition of UUI symptoms as atypical for normal ageing, with relatively less contribution of specific vascular risk factors. In contrast, severe UUI is strongly associated with previous stroke/TIA, which may result from a greater contribution of vascular disease and central nervous pathology. Patients with central lesions such as strokes and multiple sclerosis commonly report UUI as a complication. Structurally, subcortical white matter lesions, a marker of chronic vascular burden, is associated with increased symptoms of detrusor overactivity (33). However, central pathology can contribute to UUI even in the absence of overt signs of neurological disease (34): functional magnetic resonance imaging (fMRI) studies involving patients with UUI without diagnosed neurological pathology have associated UUI symptoms with abnormal activation patterns in the prefrontal and anterior cingulate cortex, insula, basal ganglia and cerebellum (34).

While this study corroborates previous findings that BMI plays a central role in the aetiology of UUI (16) (17), our results also support the emerging view that obesity contributes to UUI as part of a wider metabolic syndrome. This is shown in men but not so in women. The association between type 2 diabetes and male UUI in our study is consistent with previous studies linking UUI with markers of poor glycaemic control, including diagnosis of type 2 diabetes mellitus (35), higher HbA1c in patients with type 2 diabetes (36), raised serum insulin and increased homeostatic model assessment of insulin resistance (HOMAIR) (20). Our association of hypertension and UUI adds to a previous study linking hypertension with OAB symptoms in a female cohort (14).

This study provides prevalence of urinary incontinence subtypes for men and women within a representative large cohort in their late 60s. We demonstrate that severe UUI is a distinct disease entity while those with milder UUI may represent a broad range of contributory mechanisms. Our findings suggest that multifactorial mechanisms lead to UUI and that vascular risk factors are additionally associated, potentially playing a role in the development of pathological bladder overactivity.

## Acknowledgments

The authors thank all study members of NSHD and NSHD scientific and data collection teams. Data used in this publication are available upon request to the MRC National Survey of Health and Development Data Sharing Committee. Further details can be found at: http://www.nshd.mrc.ac.uk/data. doi: 10.5522/NSHD/Q101; doi: 10.5522/NSHD/Q102; doi: 10.5522/NSHD/Q103.

---

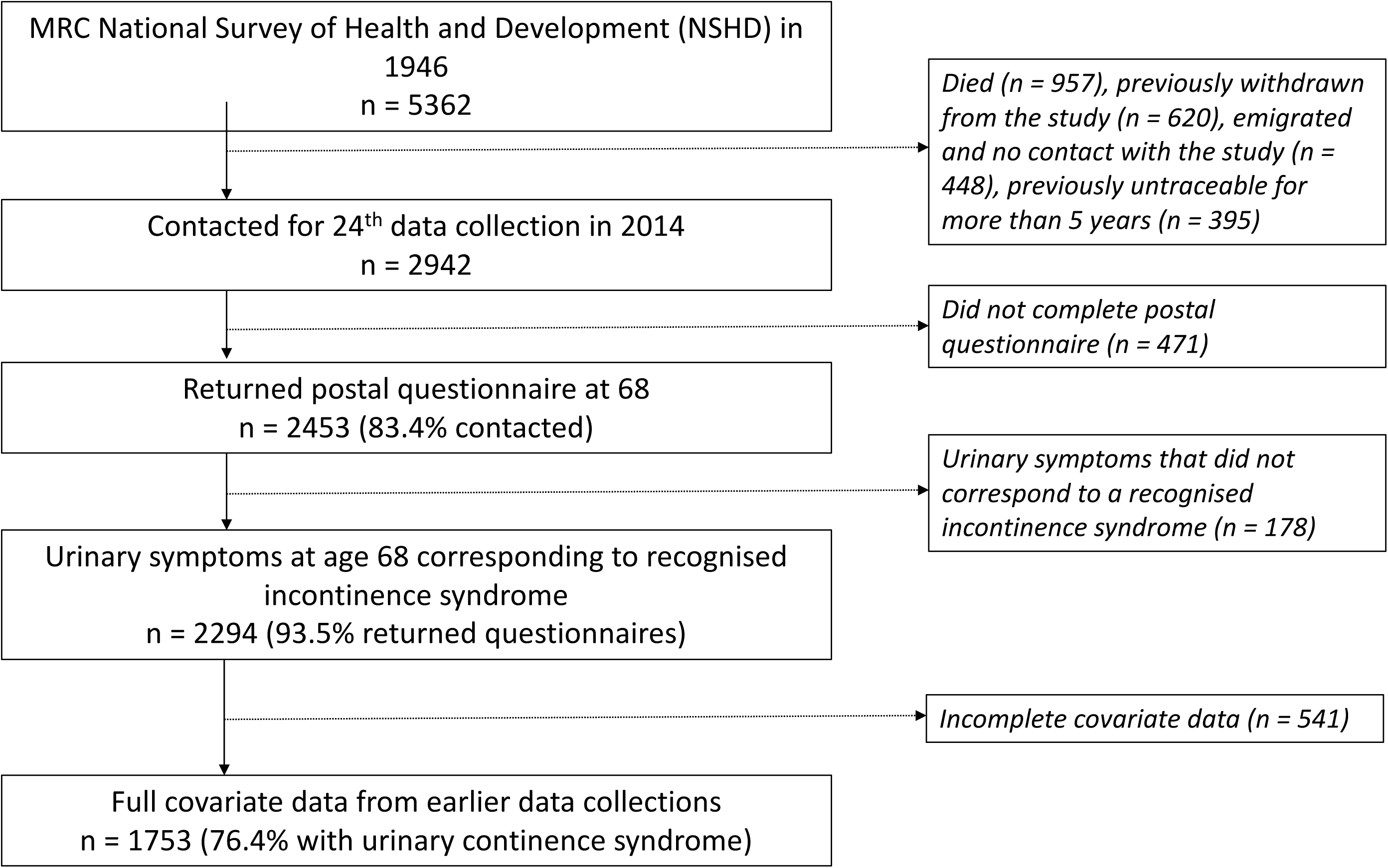

